# CircSSNN: circRNA-binding site prediction via sequence self-attention neural networks with pre-normalization

**DOI:** 10.1101/2023.02.07.527436

**Authors:** Chao Cao, Shuhong Yang, Mengli Li, Chungui Li

**Author notes:** (SY), (CL).

## Abstract

Circular RNAs (circRNAs) play a significant role in some diseases by acting as transcription templates. Therefore, analyzing the interaction mechanism between circRNA and RNA-binding proteins (RBPs) has far-reaching implications for the prevention and treatment of diseases. Existing models for circRNA-RBP identification most adopt CNN, RNN, or their variants as feature extractors. Most of them have drawbacks such as poor parallelism, insufficient stability, and inability to capture long-term dependence. To address these issues, we designed a Seq_transformer module to extract deep semantic features and then propose a CircRNA-RBP identification model based on Sequence Self-attention with Pre-normalization. We test it on 37 circRNA datasets and 31 linear RNA datasets using the same set of hyperparameters, and the overall performance of the proposed model is highly competitive and, in some cases, significantly out-performs state-of-the-art methods. The experimental results indicate that the proposed model is scalable, transformable, and can be applied to a wide range of applications without the need for task-oriented fine-tuning of parameters. The code is available at https://github.com/cc646201081/CircSSNN.

**Author summary:** In this paper, we propose a new method completely using the self-attention mechanism to capture deep semantic features of RNA sequences. On this basis, we construct a CircSSNN model for the cirRNA-RBP identification. The proposed model constructs a feature scheme by fusing circRNA sequence representations with statistical distributions, static local context, and dynamic global context. With a stable and efficient network architecture, the distance between any two positions in a sequence is reduced to a constant, so CircSSNN can quickly capture the long-term dependence and extract the deep semantic features. Experiments on 37 circRNA datasets show that the proposed model has overall advantages in stability, parallelism, and prediction performance. Keeping the network structure and hyperparameters unchanged, we directly apply CircSSNN to linRNA datasets. The favorable results show that CircSSNN can be transformed simply and efficiently without task-oriented tuning. In conclusion, CircSSNN can serve as an appealing circRNA-RBP identification tool with good identification performance, excellent scalability, and wide application scope, which is expected to reduce the professional threshold required for hyperparameter tuning in bioinformatics analysis.

## Introduction

Circular RNA (or circRNA) is a single-stranded RNA with a closed-loop structure[1, 2]. It is resistant to exonuclease-mediated degradation, and is more stable than most linear RNA. Recent studies have shown that circRNA molecules are rich in microRNA (miRNA) binding sites, which act as miRNA sponge (miRNA sponge) in cells[3–5], thus relieving the repressive effect of miRNA on its target genes and increasing the expression level of target genes. This mechanism of action is known as competitive endogenous RNA (ceRNA) mechanism. By interacting with disease-associated miRNAs, circRNA play a significant role in disease[6–8]. It has been shown that circRNA is conducive to the suppression of cancer by binding to some RBPs [9]. Therefore, an in-depth analysis of the interaction between circRNAs and RBPs to understand the development of tumor biology has remarkable significance.

Benefiting from the High-throughput sequencing of RNA isolated by crosslinking immunoprecipitation (HITS-CLIP, also known as CLIP-Seq) sequencing technology, researchers have found there are several RBP binding sites in circRNA in eukaryotes[10, 11]. Therefore, many bioinformatic methods have been proposed to predict circRNA-RBP interaction. For example, inspired by the extraction of image feature, Wang et al. proposed a circRNA-RBP classification model based on CNN, which use the RBP binding sites on CS-circRNAs to predict its relevance to cancer[12]. Based on the capsule network, the CircRB[13] model also utilized convolutional operations to extract the feature of circRNAs, and leveraged the dynamic routing algorithm to classify the binding sites. To introduce temporal information in circRNA-protein binding site, Ju et al. first used CNN to extract features, then combined LSTM with conditional random fields and proposed a sequence-tagged deep learning model to identify circRNA-protein binding sites [14].Simiarly, ZHANG et al. combined CNN and BiLSTM into a hybrid neural network in CRIP model[15]. They also use CNN to extract features and use BiLSTM to capture the temporal information and obtain long-term association information. Unlike the methods mentioned above, CRIP used a codon-based scheme to encode RNA sequences[15]. Also based on a hybrid deep network composed of CNN and BiLSTM network, Jia et al. applied XGBoost with incremental feature selection to conduct feature encoding and proposed PASSION[16] algorithm for circRNA-protein binding site prediction. Drawing on the ideas of NLP, Yang et al. proposed a KNFP (K-tuple Nucleotide Frequency Pattern) encoding scheme to describe local information, and applied word2vec to obtain global statistical information. The network architecture in Yang’s model is a hybrid model consisting of a multi-scale residual CNN and a GRU network [17]. Li et al. introduced multi-view subspace learning and ensemble neural network into Yang’s model, and proposed two models named as DMSK[18] and CRBPDL[19], respectively. The models mentioned above have made impressive improvements in performance of circRNA-RBP prediction, but there are still limitations in the description of global relation. The reason is that these methods fail to make full use of the contextual information of circRNA sequences.

To overcome this issue, inspired by the newly proposed BERT(Bidirectional Encoder Representations from Transformers) model, Yang et.al first pre-trained a DNABERT model[20], then fine-tuned DNABERT to capture the semantic and syntactic information of the initial RNA sequence, and finally used the deep temporal convolutional network(DTCN) to predict the circRNA-protein binding site[21]. However, though the existing models have tried many attempts, from single-view to multi-view, to enrich the diversity of features, they mainly resort to CNN and RNN or a hybrid of them to extract the deep feature of circRNA, there is still large rooms for improvements regarding the issues such as the poor parallelism of network architecture, inability to flexibly capture long-term dependencies of features, and insufficient algorithm stability.

In this study, we developed a novel end-to-end circRNA-binding site prediction model called CircSSNN (CircRNA-binding site prediction via Sequence Self-attention Neural Network). To capture the hierarchical relationship between nucleotide sequences, we extract the initial features of circRNA sequence by aggregating multiple gene encoding scheme, including static local context and dynamic global context information. Then, we use Transformer to design a network architecture i.e., Seq_Transformer, to extract the latent nucleotide dependencies to complete the task of CircRNA-RBP site prediction.

In the proposed model, the ResNet and LayerNorm modules are incorporated into deep network to improve the robustness and reduce the sensitivity to hyperparameters, which also allows the algorithm to generalize well to different RNA-RBP combination recognition tasks. We compared CircSSNN with several state-of-the-art baselines on 37 popular circRNA benchmark datasets to verify its effectiveness and generalization. Moreover, keeping the network structure and hyperparameter unchanged, we directly applied CircSSNN to 31 linear RNAs dataset, and also obtained better performance than existing methods. The experimental results show that CircSSNN is superior to existing methods regarding to the recognition performance, and the generalization to different types of RNA-RBP. Therefore, it can serve as a competing candidate for the task of RNA-RBP prediction with a wide range of applications.

## Materials and methods

### Datasets

To verify the effectiveness of the CircSSNN, we adopt 37 circRNA datasets as benchmark datasets following the baselines we compared[15, 16]. We first downloaded the datasets from circRNA interactome database (https://circinteractome.nia.nih.gov/). Subsequently, we obtained 335976 positive samples and 335976 negative samples following the process of iCircRBP-DHN[17].

To demonstrate the generalization of CircSSNN regarding different types of RNA-RBP, we also test the algorithm on 31 linear RNA datasets^[22, 23]^ coming from CLIP-Seq data. Each linear RNA dataset has 5000 training samples and 1000 test samples [16].

### Feature Multi-descriptors

In CircSSNN, all CircRNA fragments were encoded into three types of quantified features: KNFP for expressing different levels of local contextual features, CircRNA2Vec for capturing contextual features representing long-term dependencies, and DNABERT for describing the global embedding features with learnable position encoding.

### K-tuple nucleotide frequency pattern

To describe the local dependencies of circRNA sequences, KNFP is used to count the word frequency of substrings of circRNA with different lengths, thus the local context of different lengths can be effectively captured[24].

Fig 1 shows KNFP used in this paper consisting of three parts[17]: mononucleotide composition, dinucleotide composition and trinucleotide composition. Considering a circRNA sequence with length *n*,i.e., *S* = [*S*_1_, *S*_2_,… *S_n_*], in which *S_i_* ∈ {*A*,*G*,*C*,*U*}, K-tuple nt composition can be employed to encode the raw sequence, such that each vector representing an individual k-tuple nt composition pattern, and it contains 4^K^ components as following:

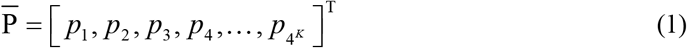

**Fig 1.**
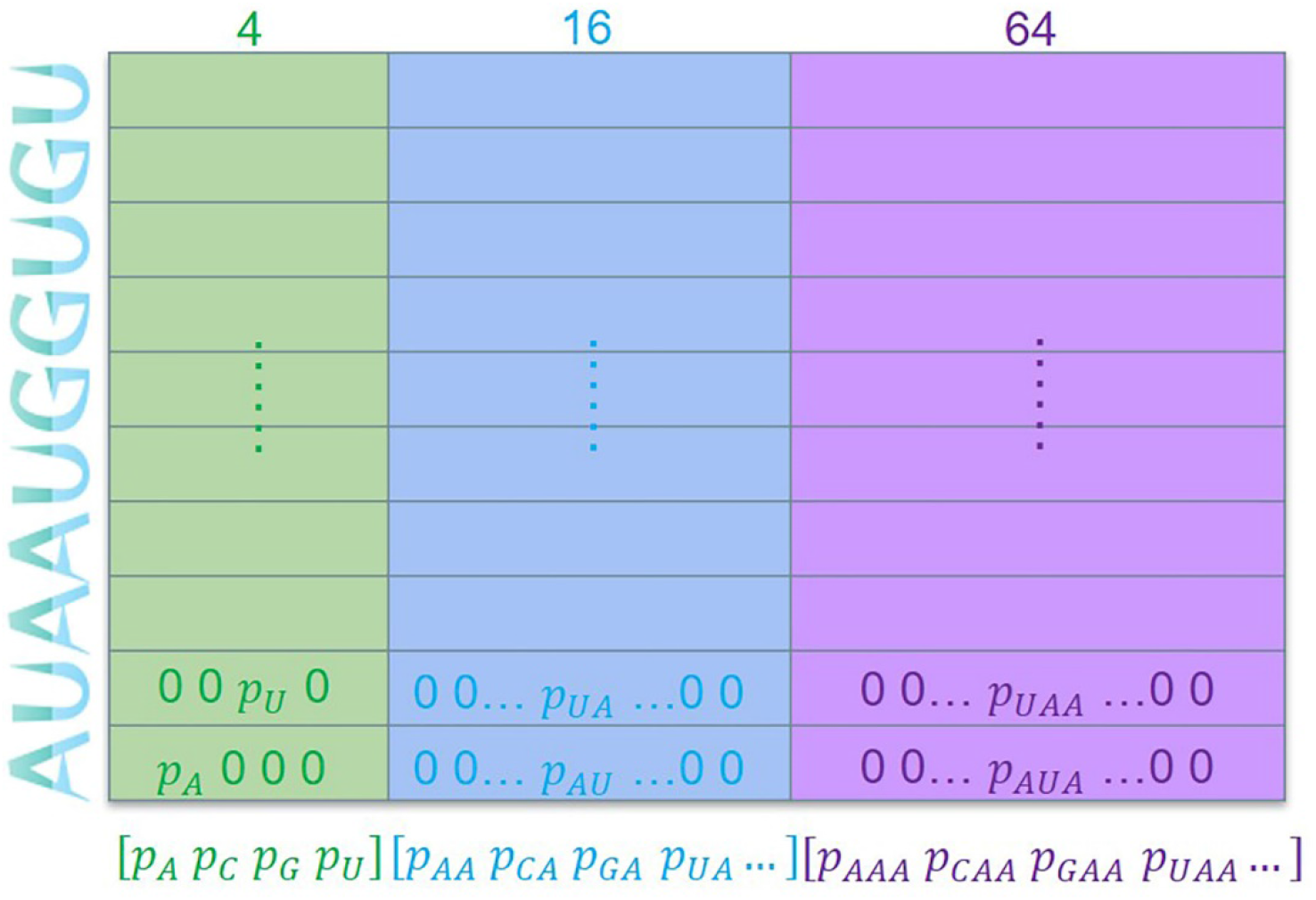
Encoding scheme of KNFP.

### CircRNA2Vec

As is shown in Fig 2, we adopt the Doc2Vec model[25] to learn the global expression of circRNAs. Doc2Vec first obtains the circRNA substrings by moving a sliding window of width 10 one letter each step over the CircRNA sequence, and then tokenize the obtained substrings into circRNA words by using the Circrna corpus from circBase[26].

**Fig 2.**
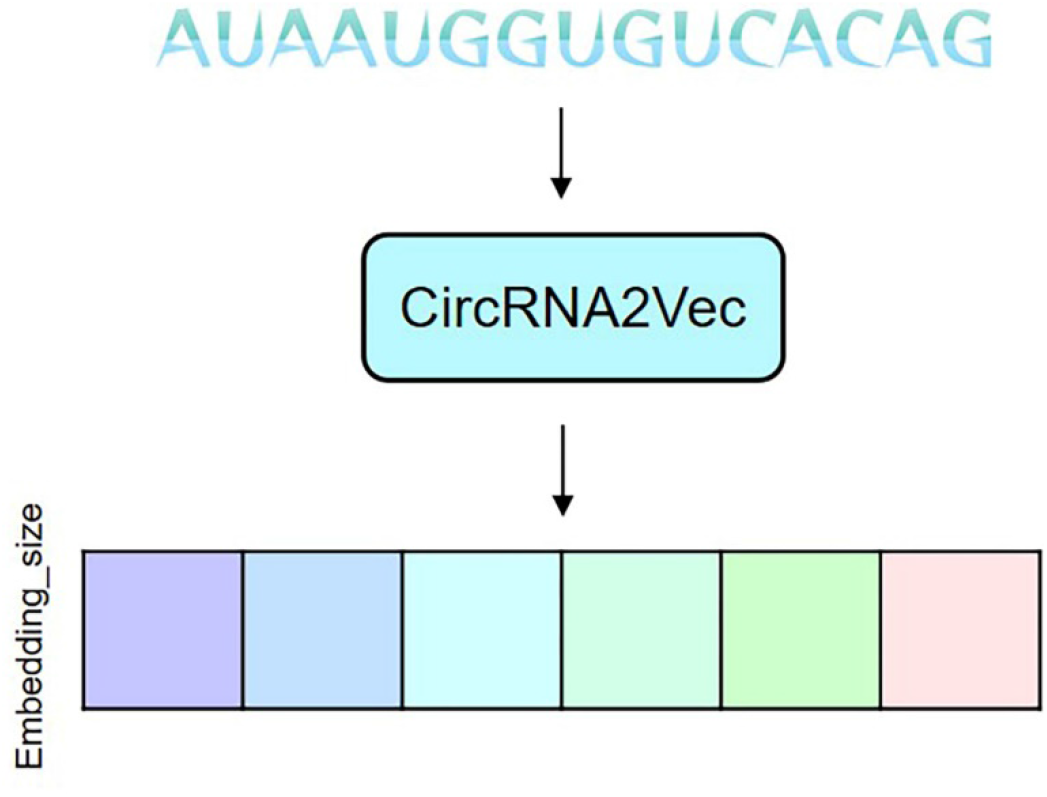
Encoding scheme of CircRNA2Vec.

We use Doc2Vec to learn the distributed expression of circRNA after tokenization. Specifically, for a central word *w_t_* obtained by tokenization, considering its context words *w*_*t*–*k*_~*w*_*t*+*k*_, the conditional probability of this central word can be modeled as following,

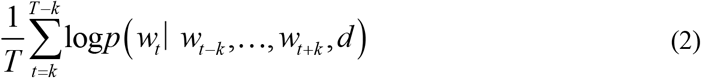

where d is the matrix of the document containing the substring considered, this is the difference between Doc2Vec and word2vec[27], i.e., the former considers the information of the document [25].

### Global embedding features based on CircRNA sequences

BERT is a language model that has achieved great success recently. Based on Transformer, BERT trains its network by unsupervised learning. Different from word2vec and Doc2Vec, BERT contains learnable positional parameters and thus can express relative position in the context. Pre-training with BERT can obtain well-generalized base parameters, which can be applied to a specific task just with corresponding fine-tuning.

Similar to HCRNet[21], we first tokenize a circRNA sequence by k-mer in which k is set as 3. Next, we perform fine-tuning on a large amount of circRNA data. Similar to the original BERT, this pre-training and fine-tuning strategy can save a lot of training time and facilitate the following learning tasks remarkably.

### Deep neural network architecture

In this section, we propose the CircSSNN framework to fully exploit the latent representation of features and facilitate the subsequent classification tasks. The overall framework is shown in Fig 3. CircSSNN consists of two parts in total, i.e., the feature encoder module and the Sequence Self-Attention Mechanism module. As stated above, multiple initial features are extracted from the raw sequence by KNFP, CircRNA2vec and DNABERT, and these initial features are first input into the feature encoder module to obtain the unified feature sequences, which are subsequently input into the next module to extract features with self-attention. The final step of classification is carried out by SoftMax.

**Fig 3.**
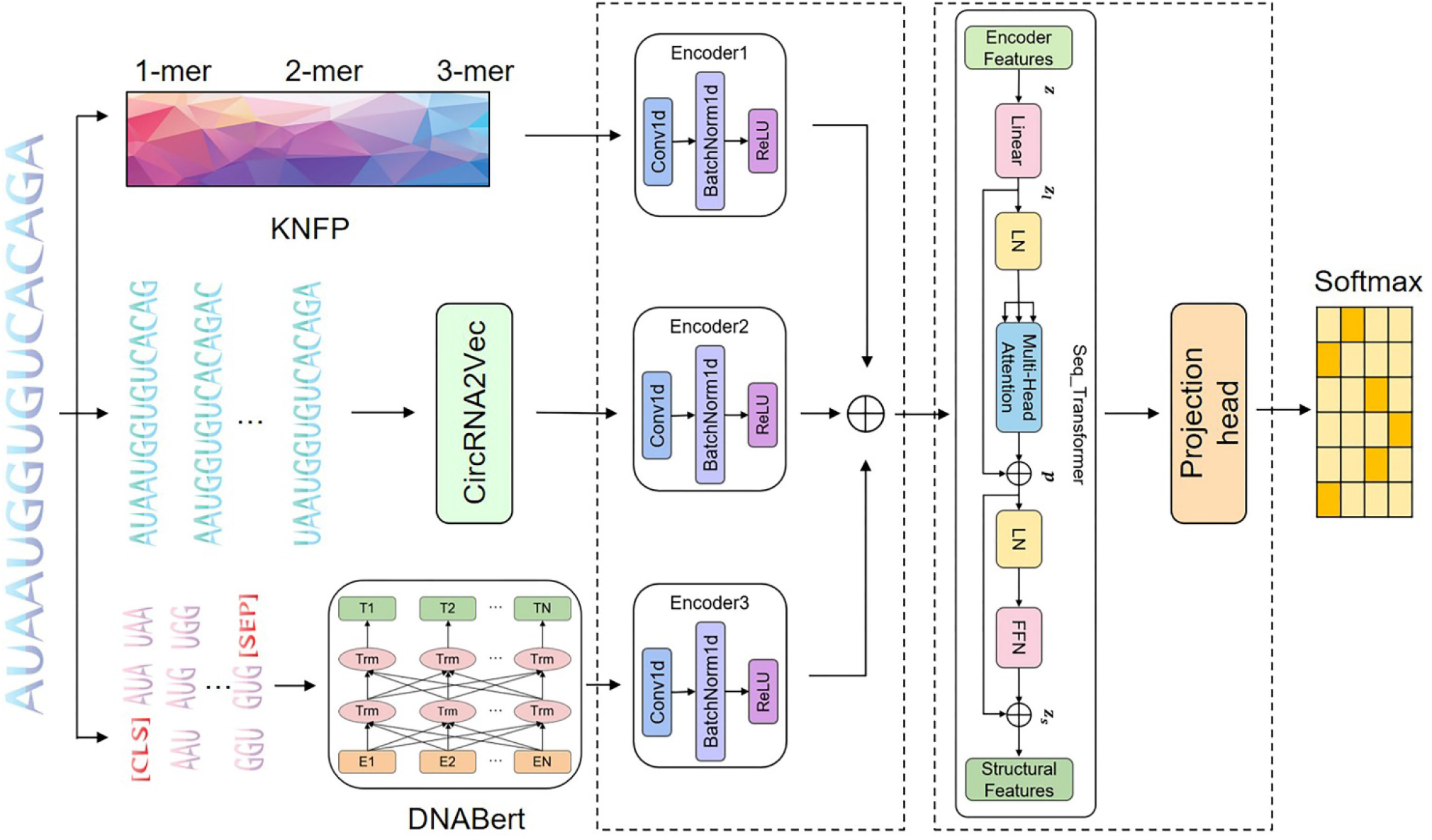
framework of CircSSNN.

### Feature encoding module

The multiple initial features obtained from different feature descriptors have inconsistent channel numbers, magnitudes, and magnitude units, etc. Such issues will hinder the later analysis. To overcome these issues, data unifying is needed to make the initial features share the same form to facilitate the subsequent feature fusion.

We construct the feature encoder layer by CNN to unify the channels of multiple initial features and conduct data normalization. As shown in Fig 4, the feature encoder layer consists of three sublayers, i.e., the one-dimensional CNN layer, the one-dimensional BatchNorm layer, and the ReLU activation function.

**Fig 4.**
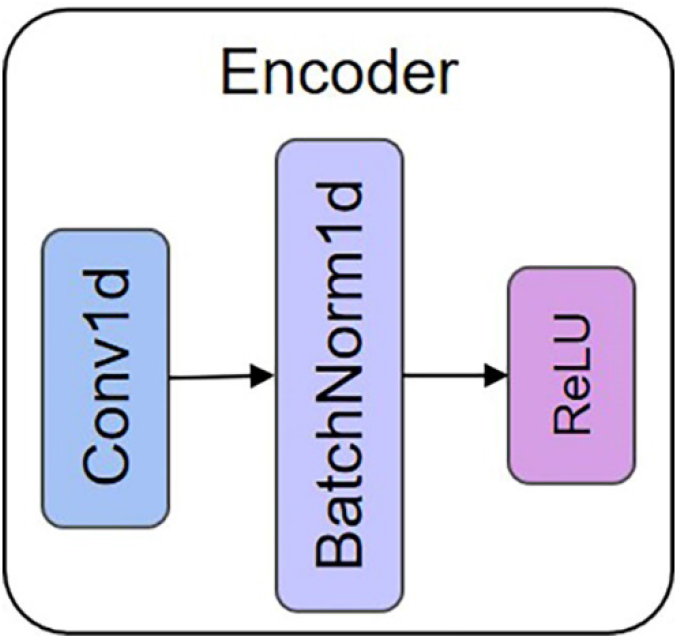
Feature encoding module.

### Sequence Self-Attention Mechanism Module

Transformer[28] is a network architecture based on attention mechanism and abandoned traditional CNN and RNN. More precisely, Transformer consists only of Self-Attention and Feedforward Neural Network (FNN). This simple architecture of Transformer brings better performance, higher parallelism, and less time-complexity. It has been successfully applied to various fields such as NLP and CV, and many researchers[29–31] have incorporated Transformer as a sub-model and achieved impressive success.

We partially adopt the architecture of Transformer with slight modification as the extractor of deep structure, i.e., the Seq_Transformer as shown in Fig 5.

**Fig 5.**
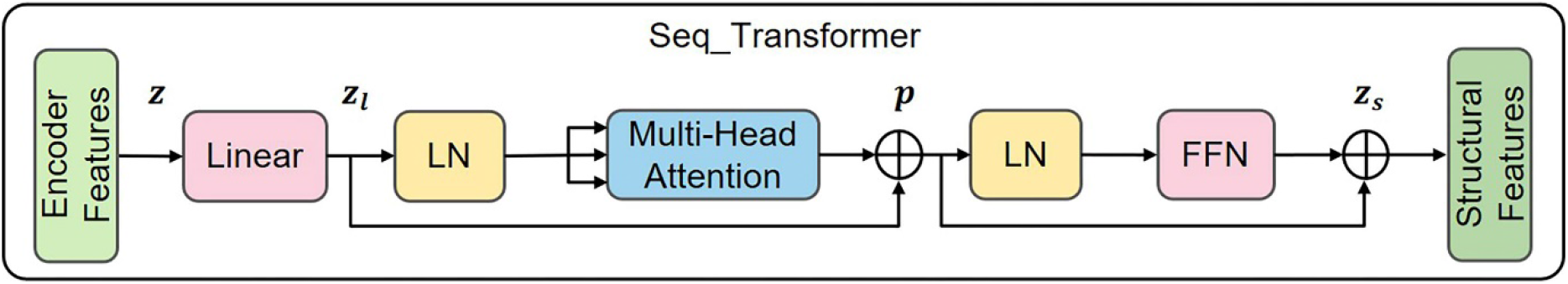
the structure of Seq_Transformer.

When constructing a neural network using the Transformer architecture superimposing multiple sub-layers, either in the encoder or in the decoder, leads to poor information propagation through the network, thus making the training very difficult.[32, 33] To overcome this issue, we leverage the residual module to improve the efficiency of information propagation and conduct layer normalization to reduce the variance of the sub-layers. There are two ways to incorporate layer normalization into the residual network. Let *F* be a sub-layer (either in the encoder or decoder) in the Transformer architecture, and denote its parameter set by *θ_l_*

#### Post-Norm

In the pioneering works of Transformer[28], it is common practice to do residual addition followed by Layer Normalization(LN) as follows,

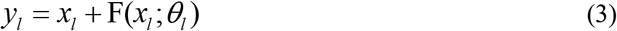

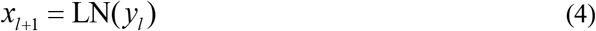

#### Pre-Norm

In recent years, many researchers[34] prefer to conduct Layer Normalization(LN) on the inputs of sublayers rather than the outputs, like this,

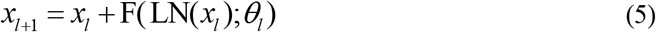

The effect of Post-Norm or pre-Norm is comparable for shallow networks. Both methods can effectively improve the distribution of parameters, which facilitates the smooth training. However, for a deeper network, it is has been pointed out that Pre-norm is better than Post-norm [32, 33]. Specifically, for CircSSNN, since DNABERT is used in the initial feature extraction and the Seq_Transfomer is designed next, the network is rather deep in general. Therefore, for the cirRNA-RBP prediction, which is task of the proposed model, we argue that the Pre-norm is more effective than the Post-norm. We have empirically demonstrated this point in the ablation experiments in the Section of Results and discussion.

Theoretically, this phenomenon can be explained by a carefully examining of the nature of network training. It is well known that the training network is essentially the backward propagation of error computed by the loss function and the correspondent adjustment of weight parameters of the network according to the error propagation. Take a submodule containing L-layers for example, the error back-propagated from the next layer is represented by *ε*, and *x_L_*, represents the output of the last layer. If the Transformer adopts the Post-Norm strategy, according to the chain rule, the partial derivative of *ε* with respect to *x_L_*, can be calculated for a particular sublayer *x_l_* as follows[33],

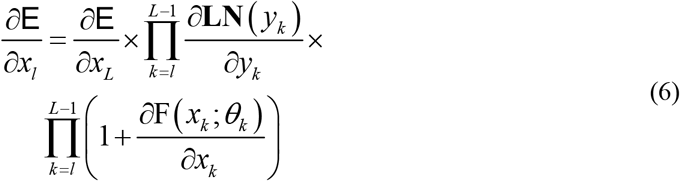

where 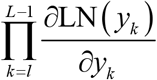 denotes the normalized information which is propagated backward, and 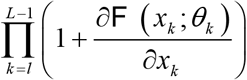 indicates the information is back-propagated through the residual module. Similarly, for the case of Pre-norm, we can obtain the gradient as follows[33],

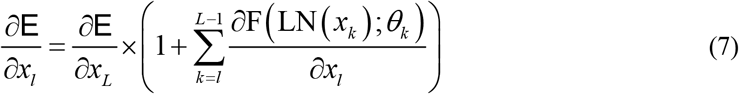

From Eq.(7), it is easy to find out that the term “1” in the parenthesis enable the direct backward propagation of 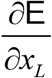 from the last layer to the *l*-th layer, i.e., the propagation through the residual module no longer depend on the number of layers.

Comparing the calculation of the information propagation of residual module in Eq. (6) and Eq. (7), one can find that in Eq. (6) the information passing through residual module do not propagate directly from layer L to layer *l*. This is because in Post-norm, the residual connection module are not a real bypass of layer-normalization layer, resulting in a concatenated multiplicative term for the gradient propagation of the residual module in Eq. (6), i.e., 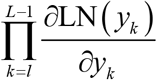, in which it can be found obviously, if the number of layers goes deeper, this term will suffer from gradient vanishing or exploding.

Therefore, our model is connected by Pre-norm residual blocks[32, 33], and features are normalized before passing through the multi-headed self-attention network, thus producing a more stable gradient.

The overall process of CircSSNN is as follows. We first extract multiple initial features by KNFP, CircRNA2vec, and DNABERT,. These initial features are then integrated into multi-view fused feature *z_l_* which is divided into two ways using the residual connection module as follows,

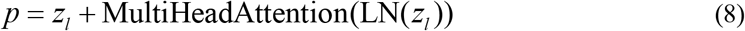

In Eq.(8), one way of information remained as it was and propagated from right to left directly, while the other way of information was first normalized by Pre-norm LN before passing through the MHA module. The Pre-norm LN is defined as,

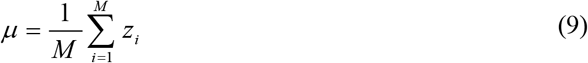

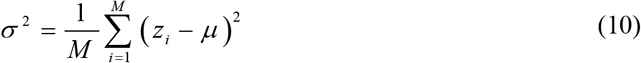

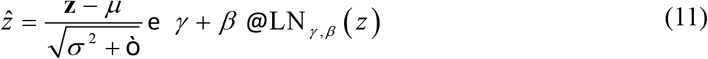

In Eqs(9–11), *M* is the number of neurons. Features are extracted using scaled dot-product multi-head attention to capture contextual feature as follows,

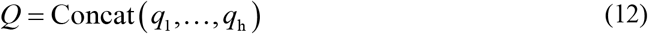

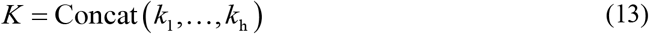

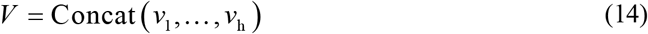

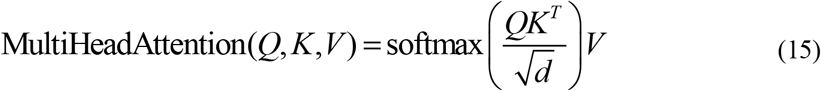

In Eqs(12–14), *h* is the number of heads, *q_i_*,*k_i_* and *v_i_*, *i* ∈ {1,2,…*h*} denote the query, key, and value respectively. *Q,K* and *V* indicate the aggregation of multiple *q_i_*, *k_i_*, and *v_i_*, respectively. In Eq.(15), *d* is the dimension of the input vector. Then, the information passing through the MHA module and bypassing it are added together to get *p* as described in Eq.(8). Similarly, before the information passing through the FFN module, it is also processed by Prenorm LN. By this way, the input information is finally turned into a unified structured deep feature to conduct the subsequent classification.

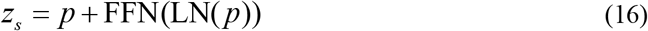

From the network architecture of CircSSNN, one can find it differs from the existing models in two aspects.

First, to the best of our knowledge, this is the first attempt to introduce residual module with Pre-norm LN in CircRNA recognition. As stated in [32, 33], the residual module with Post-norm LN brought about higher risk of gradient vanishing or exploding when the network goes deeper. Therefore, we adopt the Pre-norm LN scheme to avoid this problem while using residual connection to improve the efficiency of information transmission.

Second, we proposed the Seq_Transformer module based on the self-attention to extract temporal contextual features. Most of existing works proposed for CircRNA-RBP prediction, such as DMSK[18], CRBPDL[19], iCircRBP-DHN[17], and CRIP, etc., mainly use RNN such as LSTM or GRU for capturing temporal dependence. However, the computation of RNN or its variants is sequential, i.e., calculatiing results of time step *t* must depend on that of time step *t*-1, which dramatically limits the parallelism. In addition, long-term dependency is prone to loss during propagation along the sequential RNN network. LSTM and GRU adopted some gating mechanisms to mitigate this problem to a certain extent, but the effectiveness of gating mechanisms is undesirable for long-term dependencies. Therefore, compared with the models based on the Self-Attention mechanism, these models suffer from insufficient parallelism and poor ability to capture long dependencies. However, Attention mechanism has seldom been employed to extract features directly in this field. Up to now, only Yang et al. used the Attention mechanism in the iCircRBP-DHN model they proposed in 2020. But in iCircRBP-DHN, Attention mechanism was not employed as a direct feature extractor but as a supplement to the GRU mechanism, i.e., iCircRBP-DHN use Attention module to capture features after GRU processing, which to some extent destroys the dependency relationship of the original data and makes the Attention mechanism play little role. In their subsequent work, i.e., the HCRNet proposed in 2022, they has omitted the Attention mechanism. In HCRNet, Yang et al. used DTCN to extract discriminative information from hybrid features and combine the parallelism of CNN with residual connection, and thus making various perceptual field sizes available and gradients stable. DTCN alleviates the limitation of RNN regarding to parallelism to some extent. However, it is still limited by the fixed perceptual field size of CNN, and the two issues of existing models, i.e., insufficient parallelism and inefficiency in capturing long-term dependencies, still exist. In contrast, in CircSSNN, after the initial multiple features were integrated into a unified one, feature extraction is performed directly using Seq_Transformer without intermediate processing by RNN or its variants. As a result, we solved the above two issues by adopting the Seq_Transformer. The advantages of Seq_Transformer can be analyzed as follows. First, it is constructed based on the Attention mechanism rather than sequential structure, therefore its calculation can be performed in the format of matrix multiplication, which can be well parallelized and accelerated by modern deep learning framework based on GPU. Second, by using Seq_Transformer, the distance between any two positions in the sequence can be reduced to a constant, and long-term dependence can be effectively captured. In addition, due to the excellent parallelism of Seq_Transformer, we can make the full use of multi-headed attention to focus on contextual information from different locations simultaneously. Therefore, the deep structure features extracted by Seq_Transformer have good classification performance.

## Results and discussion

### Experimental setting

For both circRNA and linRNA datasets, 80% of the samples were randomly selected as training data. The remaining 20% of them were used as test data. To show the generalization of CircSSNN rather than the performance improvement brought by hyperparameter tuning, we didn’t set validation sets for hyperparameter tuning in experiments. The hyperparameters of CircSSNN were set to be the same across all datasets, which eliminates the trouble of hyperparameters tuning.

We used Adam as optimizer, and set the weight_decay and batch size as 3e-4 and 64 respectively. The learning rate of Adam was controlled by the built-in learning rate scheduler of Pytorch in which the parameter *initial_rate* was set be 3e-3. As the Seq_Transformer can capture deep feature effectively and quickly, we let the learning rate decay to one-tenth every two rounds to accelerate the convergence.

### Experimental results on circRNA datasets

We compared CircSSNN with seven baselines on 37 circRNA-RBP datasets. To be fair, all the parameters were set as reported in the corresponding papers.

Four metrics include AUC, ACC, precision, and recall, were used to compare the performance of the competing methods. The performances of all methods, averaging over 37 circRNA datasets, were shown in Figure 6. The color of the scatter corresponds to the value regarding each metric, and the larger the scatter, the better the performance.

**Fig 6.**
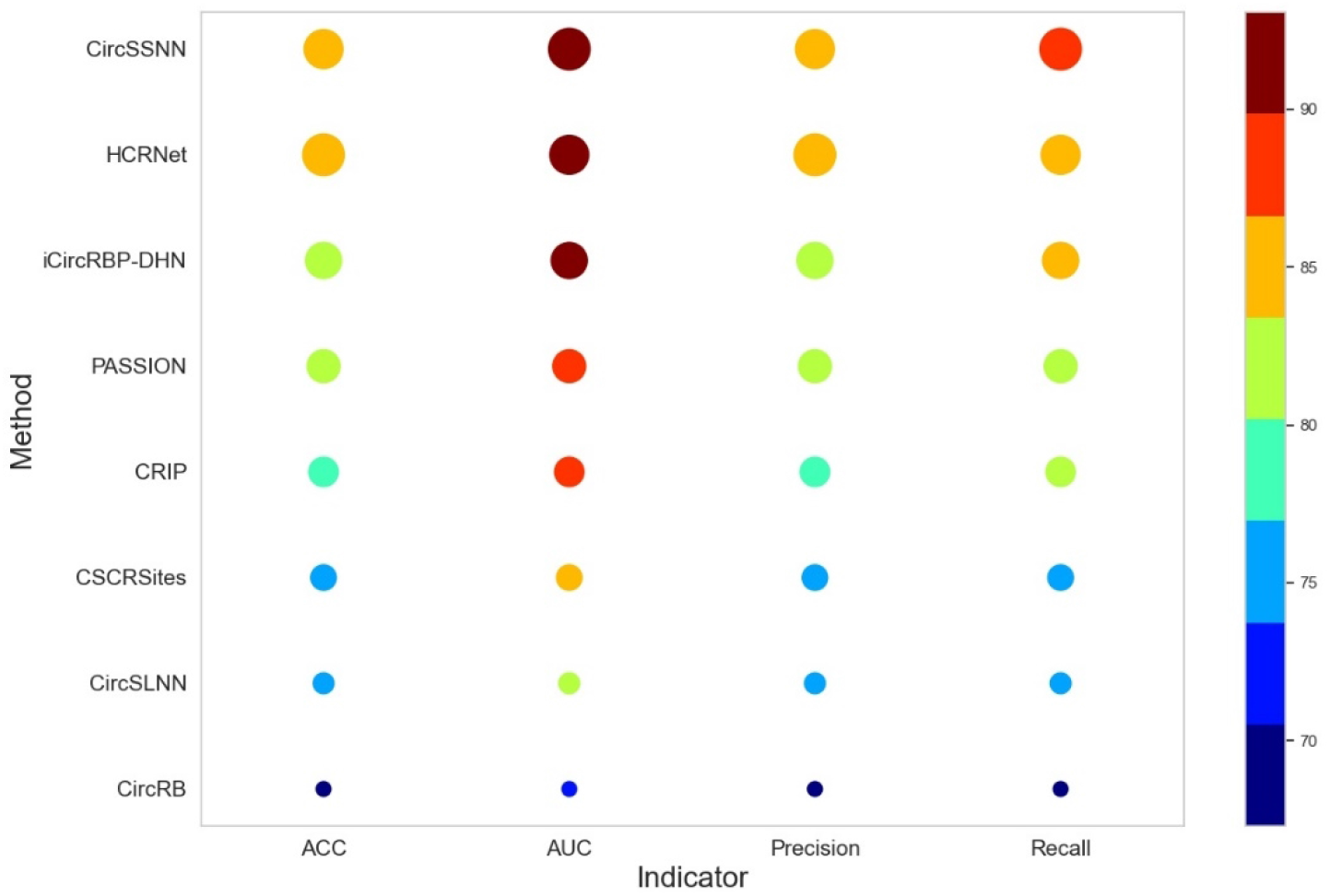
The average performance of competing methods on 37 circRNA datasets.

As can be seen from Fig 6, the performance of CircSSNN is superior to all competing methods regarding to AUC and Recall, and is slightly inferior to HCRNet regarding ACC and Precision, but is higher than the other six methods by a large margin. The detailed average value of different methods regarding ACC, AUC, Precision and recall are 85.71%, 93.07%, 85.14%, 86.69% for CircSSNN; 85.81%, 93.04%, 85.68%, 86.35% for HCRNet. As the performance of other baselines are obviously by far inferior to that of the two methods mentioned above, we don’t list them here for simplicity. The detailed AUC values are summarized in Table 1.

**Table 1.** The AUC of competing methods on 37 circRNA datasets.

| dataset | CircSSNN | HCRNet | iCircRBP-DHN | PASSION | CRIP | CSCRites | CircSLNN | CircRB |
| --- | --- | --- | --- | --- | --- | --- | --- | --- |
| AGO1 | **0.921** | **0.930** | 0.898 | 0.909 | 0.905 | 0.851 | 0.844 | 0.750 |
| AGO2 | **0.847** | **0.854** | 0.797 | 0.822 | 0.811 | 0.755 | 0.715 | 0.624 |
| AGO3 | **0.950** | **0.939** | 0.920 | 0.909 | 0.895 | 0.820 | 0.860 | 0.718 |
| ALKBH5 | **0.991** | **0.989** | 0.979 | 0.752 | 0.721 | 0.798 | 0.583 | 0.593 |
| AUF1 | **0.988** | **0.988** | 0.985 | 0.979 | 0.980 | 0.940 | 0.971 | 0.938 |
| C17ORF85 | 0.982 | **0.982** | **0.987** | 0.860 | 0.813 | 0.815 | 0.721 | 0.634 |
| C22ORF28 | **0.947** | **0.936** | 0.913 | 0.894 | 0.876 | 0.878 | 0.797 | 0.731 |
| CAPRIN1 | **0.912** | **0.905** | 0.858 | 0.860 | 0.843 | 0.827 | 0.745 | 0.685 |
| DGCR8 | **0.915** | 0.914 | 0.906 | **0.917** | 0.914 | 0.870 | 0.846 | 0.770 |
| EIF4A3 | **0.847** | **0.845** | 0.799 | 0.823 | 0.812 | 0.820 | 0.717 | 0.662 |
| EWSR1 | **0.955** | **0.946** | 0.942 | 0.938 | 0.936 | 0.882 | 0.906 | 0.805 |
| FMRP | **0.925** | **0.932** | 0.892 | 0.900 | 0.898 | 0.890 | 0.826 | 0.737 |
| FOX2 | **0.961** | **0.965** | 0.958 | 0.830 | 0.815 | 0.755 | 0.602 | 0.535 |
| FUS | **0.883** | **0.888** | 0.855 | 0.859 | 0.858 | 0.799 | 0.770 | 0.697 |
| FXR1 | **0.995** | 0.984 | **0.994** | 0.959 | 0.952 | 0.871 | 0.942 | 0.838 |
| FXR2 | **0.969** | **0.960** | 0.939 | 0.941 | 0.938 | 0.868 | 0.896 | 0.774 |
| HNRNPC | 0.974 | **0.981** | **0.977** | 0.976 | 0.972 | 0.973 | 0.970 | 0.941 |
| HUR | **0.908** | **0.906** | 0.867 | 0.879 | 0.874 | 0.850 | 0.796 | 0.666 |
| IGF2BP1 | **0.889** | **0.884** | 0.843 | 0.845 | 0.843 | 0.835 | 0.760 | 0.679 |
| IGF2BP2 | **0.871** | **0.879** | 0.831 | 0.827 | 0.821 | 0.752 | 0.740 | 0.644 |
| IGF2BP3 | **0.868** | **0.870** | 0.816 | 0.831 | 0.822 | 0.754 | 0.706 | 0.635 |
| LIN28A | **0.899** | **0.899** | 0.857 | 0.875 | 0.865 | 0.840 | 0.777 | 0.671 |
| LIN28B | **0.919** | **0.911** | 0.892 | 0.889 | 0.882 | 0.758 | 0.822 | 0.731 |
| METTL3 | **0.892** | **0.895** | 0.852 | 0.878 | 0.854 | 0.808 | 0.772 | 0.731 |
| MOV10 | **0.870** | **0.862** | 0.838 | 0.845 | 0.849 | 0.778 | 0.777 | 0.698 |
| PTB | **0.851** | **0.852** | 0.822 | 0.829 | 0.826 | 0.692 | 0.738 | 0.663 |
| PUM2 | **0.974** | **0.978** | 0.970 | 0.952 | 0.953 | 0.936 | 0.932 | 0.854 |
| QKI | **0.988** | **0.988** | 0.971 | 0.927 | 0.921 | 0.866 | 0.866 | 0.807 |
| SFRS1 | **0.981** | **0.978** | 0.964 | 0.965 | 0.964 | 0.963 | 0.926 | 0.836 |
| TAF15 | **0.996** | **0.996** | 0.992 | 0.967 | 0.965 | 0.941 | 0.968 | 0.883 |
| TDP43 | **0.936** | **0.940** | 0.926 | 0.934 | 0.930 | 0.923 | 0.896 | 0.829 |
| TIA1 | **0.982** | **0.972** | 0.961 | 0.935 | 0.932 | 0.915 | 0.901 | 0.827 |
| TIAL1 | **0.940** | **0.918** | 0.917 | 0.906 | 0.902 | 0.898 | 0.871 | 0.820 |
| TNRC6 | 0.953 | **0.981** | **0.967** | 0.785 | 0.741 | 0.729 | 0.662 | 0.550 |
| U2AF65 | **0.934** | **0.932** | 0.926 | 0.930 | 0.928 | 0.911 | 0.899 | 0.787 |
| WTAP | **0.973** | **0.983** | 0.967 | 0.794 | 0.793 | 0.808 | 0.732 | 0.621 |
| ZC3H7B | **0.852** | **0.864** | 0.804 | 0.804 | 0.792 | 0.794 | 0.697 | 0.634 |
| AVG | **0.931** | **0.930** | 0.908 | 0.885 | 0.876 | 0.842 | 0.809 | 0.729 |

**Note: The top-2 results of every column are highlighted in bold.**

Apparently, CircSSNN outperformed other competing baselines on 18 out of 37 circRNAs datasets, and produced the highest average AUC 93.1%. The number of samples of the 37 benchmark datasets ranges from 892 to 40000, which validated that CircSSNN is applicable for datasets with an extensive range of scales. Even for small scale datasets, CircSSNN still achieved competing performance. The results verify the Seq_transformer adopted in CircSSNN can effectively capture the semantic and global context of sequences and produce discriminative features. Although the improvement of CircSSNN over HCRNet was not very remarkable, HCRNet needed to tune its hyperparameters by validate sets, which is time-consuming and laborious. In contrast, CircSSNN used the same set of hyperparameters for all datasets, i.e., it didn’t need validate sets to fine-tune the hyperparameters, which demonstrated that CircSSNN was more flexible and insensitive to hyperparameters. This appealing characteristic made it easier to use, especially for non-computer professionals.

To demonstrate the stability of CircSSNN, we selected a moderate scale dataset TIAL1 with 10,912 samples, and repeated the test of the top two models, i.e., CircSSNN and HCRNet, ten times on TIAL1. The fluctuation of performance was illustrated in Fig 7. In Fig 7, the curve of CircSSNN fluctuated more mildly than that of the HCRNet. It further illustrated that the Seq_Transformer used in CircSSNN was more flexible, and less affected by sample randomness, and the features extracted by Seq_Transformer are more stable.

**Fig 7.**
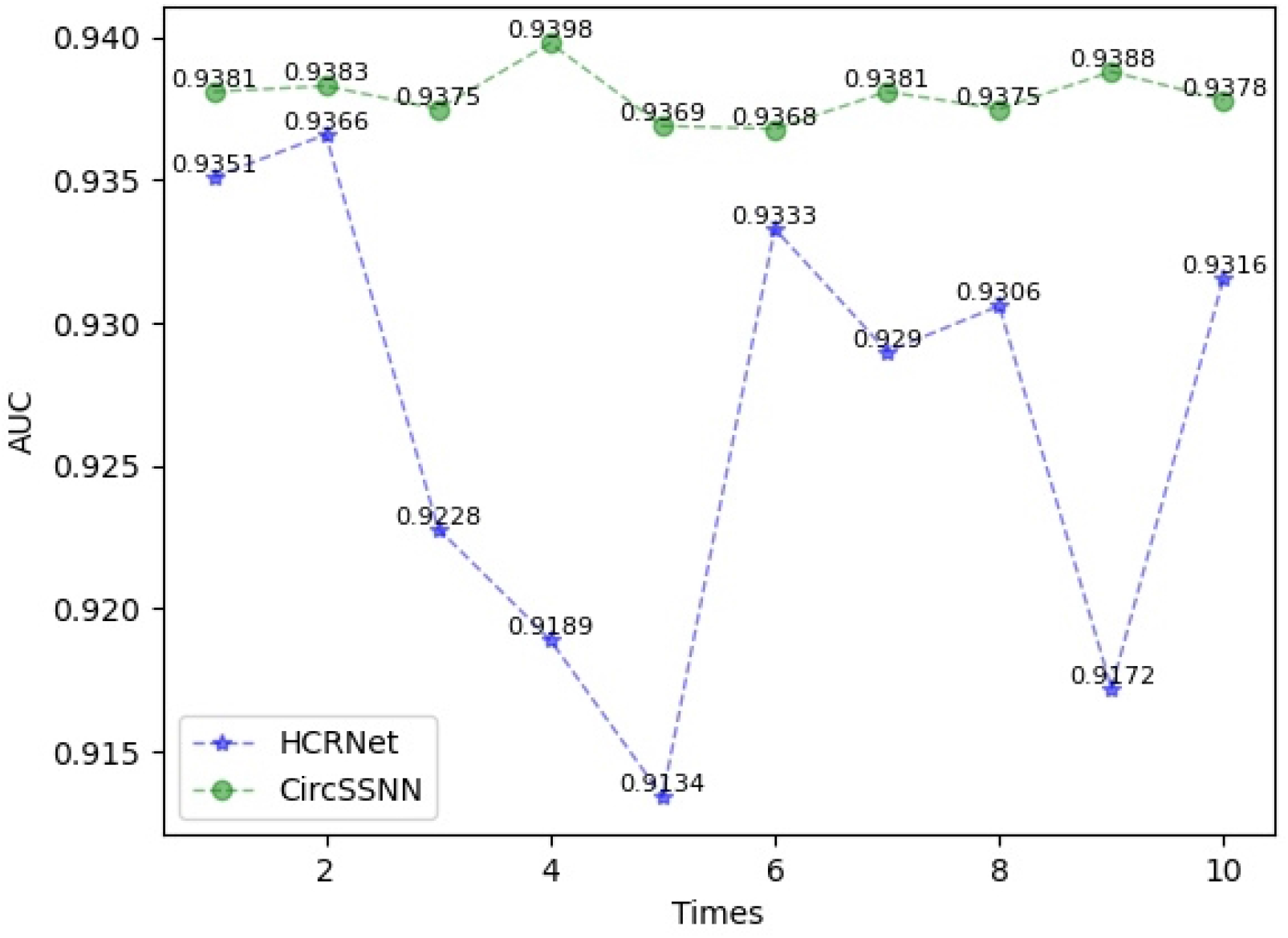
Comparison of the stability of HCRNet.

To compare the efficiency and parallelism of CircSSNN and HCRNet, we trained the two models on 37 circRNA datasets ten times with the same hardware and software configuration, and the results showed the average training times of the two models are 10 hours and 13 hours, respectively, which showed that CircSSNN was more efficient and parallelizable. The reason is that the Seq_transformer used in CircSSNN is entirely based on the Attention mechanism, which converts data into Query, Key, and Value at the same time, and thus facilitates the parallel retrieval of feature information.

To demonstrate the advantage of Pre-norm over Post-norm, we kept the other modules of the CircSSNN unchanged, and compared the effect of Pre-norm and Post-norm on 37 circRNA datasets. In Fig 8, the blue bar represents the performance of CircSSNN with Post-norm strategy, while the red bar represents the performance of Pre-norm. As shown in Fig 8, the Pre-norm strategy brings performance gains on 36 out of 37 datasets, with an increase of more than two percents on about half of the datasets.

**Fig 8.**
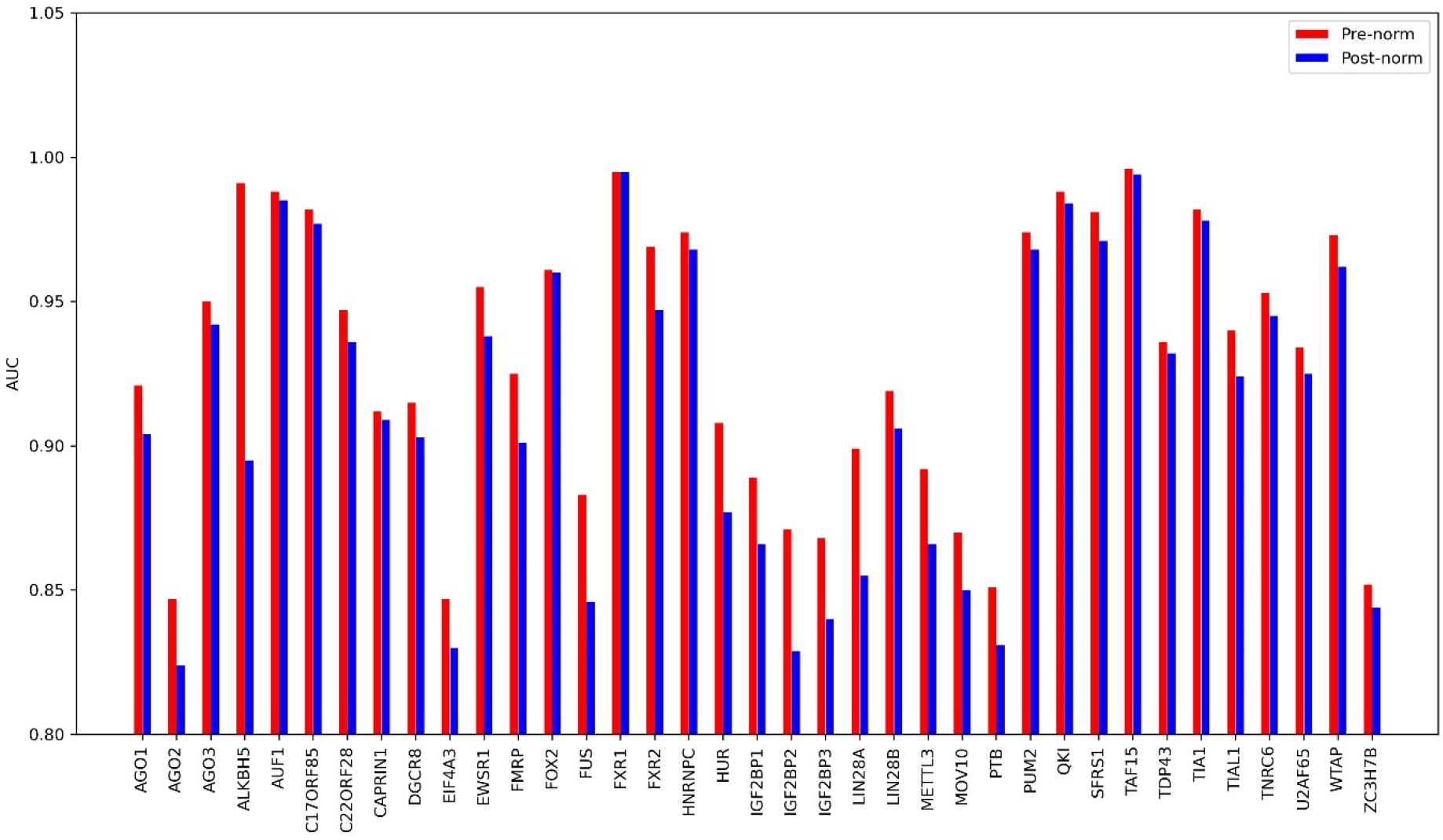
Comparison of the effect of pre-norm and post-norm on 37 circRNA datasets.

### The prediction performance of CircSSNN on linear RNA datasets

The CircSSNN is highly transformable, and can be applied to other types of RNA-RBP prediction tasks without hyperparameters tuning. To verify this, we tested CircSSNN and baselines on 31 linear RNA datasets, and the results were shown in Fig 9. As shown in Fig 9, without hyperparameters tuning, CircSSNN achieved favorable performance over other state-of-the-art baselines, which demonstrated CircSSNN was stable and transformable. The detailed value of AUC was listed in Table 2. As the models designed for cirRNA datasets, such as HCRNet and iCircRBP-DHN, do not specify the necessary detailed operation and parameter settings for migrating them from cirRNA dataset to linRNA dataset, we cannot reproduce the results of these models in our experiments, and just list in Tab.2 the AUC values published in their original papers for comparison. However, as can be observed in Fig 9 and Tab.2, even though compared with their results which was produced after fine-tuning of hyperparameters with validate sets, the results of CircSSNN, which was obtained without hyperparameters tuning, still outperformed these models in most cases. In detail, the proposed CircSSNN achieved the best AUC on 21 out of 31 linear RNA dataset, and the average value of AUC is 0.931, which is 0.7 percent higher than that of HCRNet. In some dataset, CircSSNN outperforms HCRNet by quite a bit margin, for example, AUC of CircSSNN is 4.5 and 3.6 percent higher than HCRNet on hnRNPL 1 and hnRNPL-2, respectively. Therefore, even directly keeping unchanged the network architecture and parameters designed for circRNA, CircSSNN can still produce competitive results when applied to linear RNA datasets.

**Fig 9.**
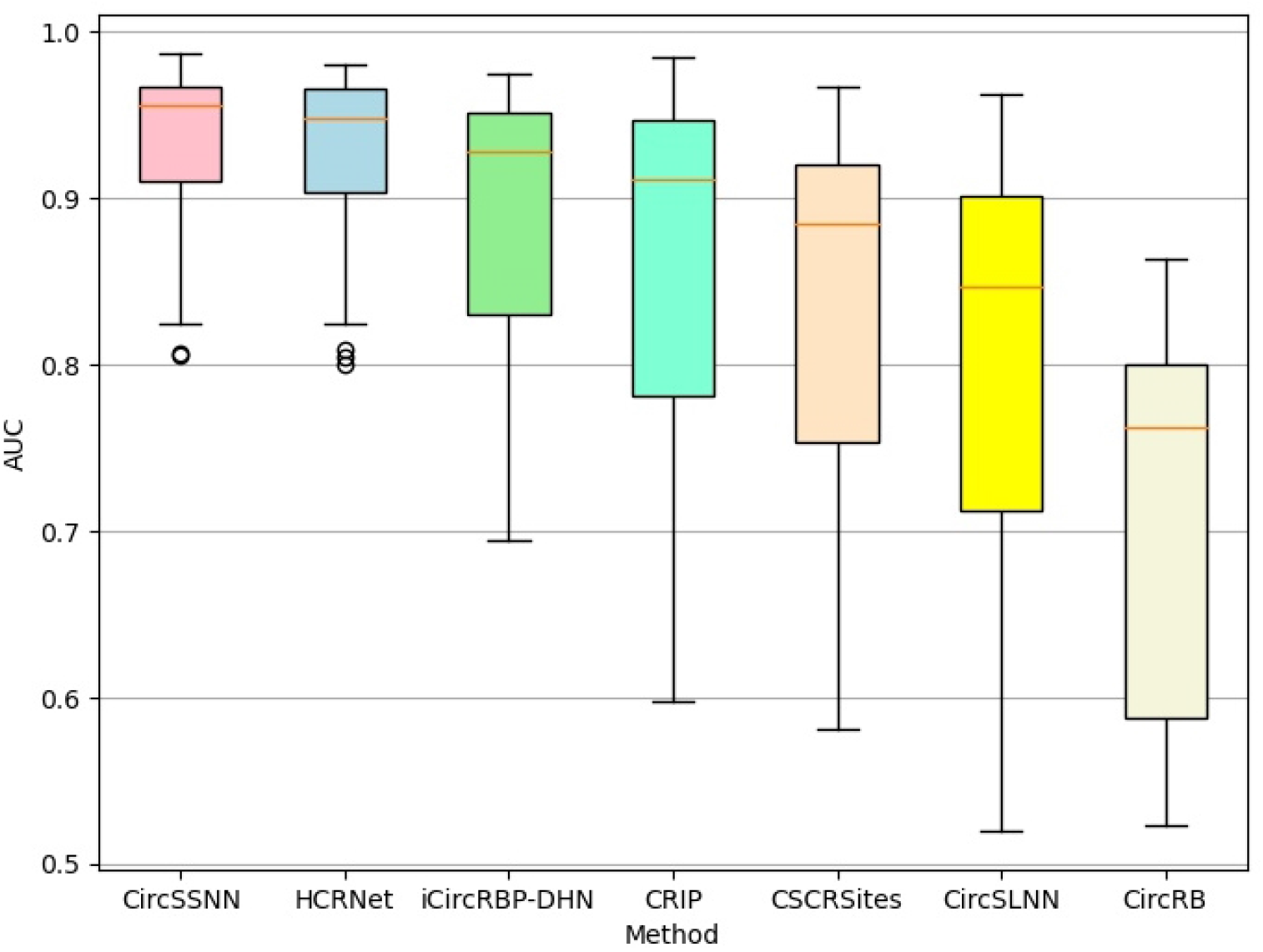
Boxplot comparison results of different models on 31 linear RNA datasets regarding to AUC.

**Table 2.**
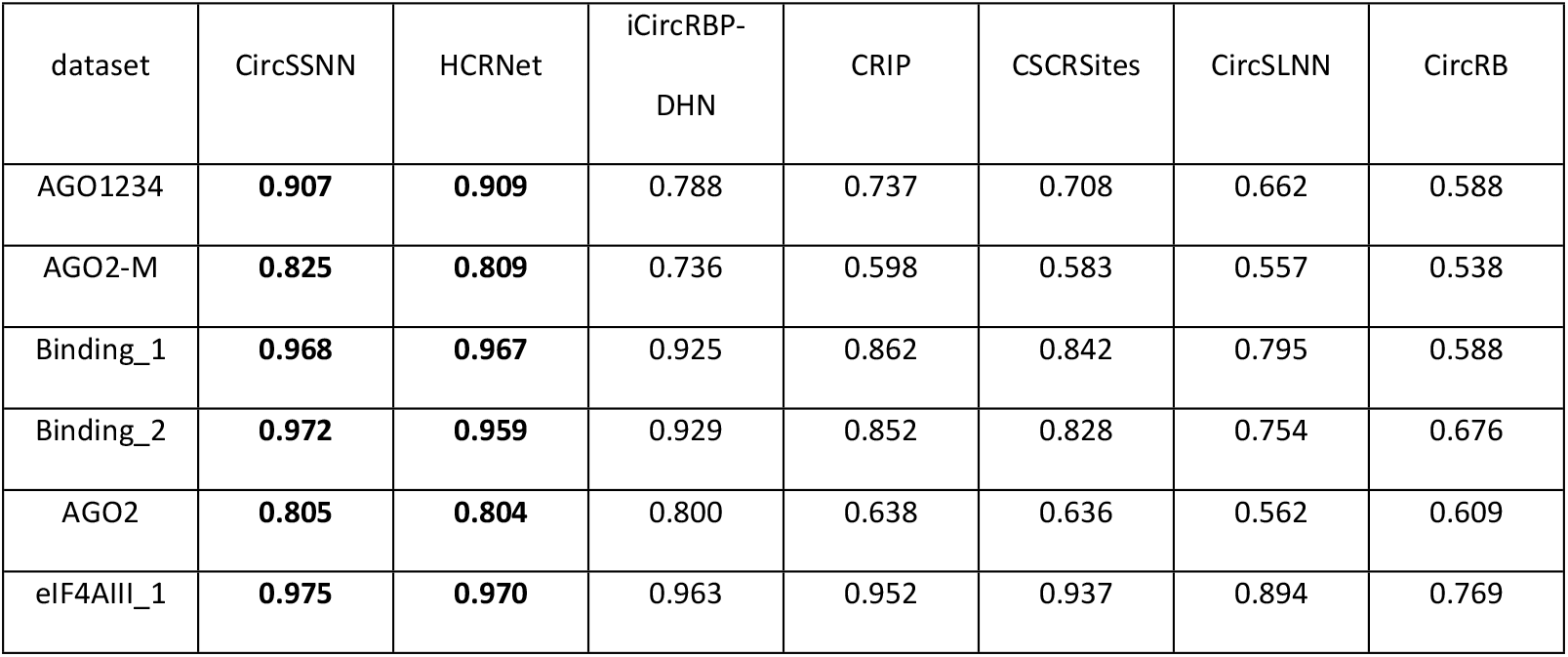
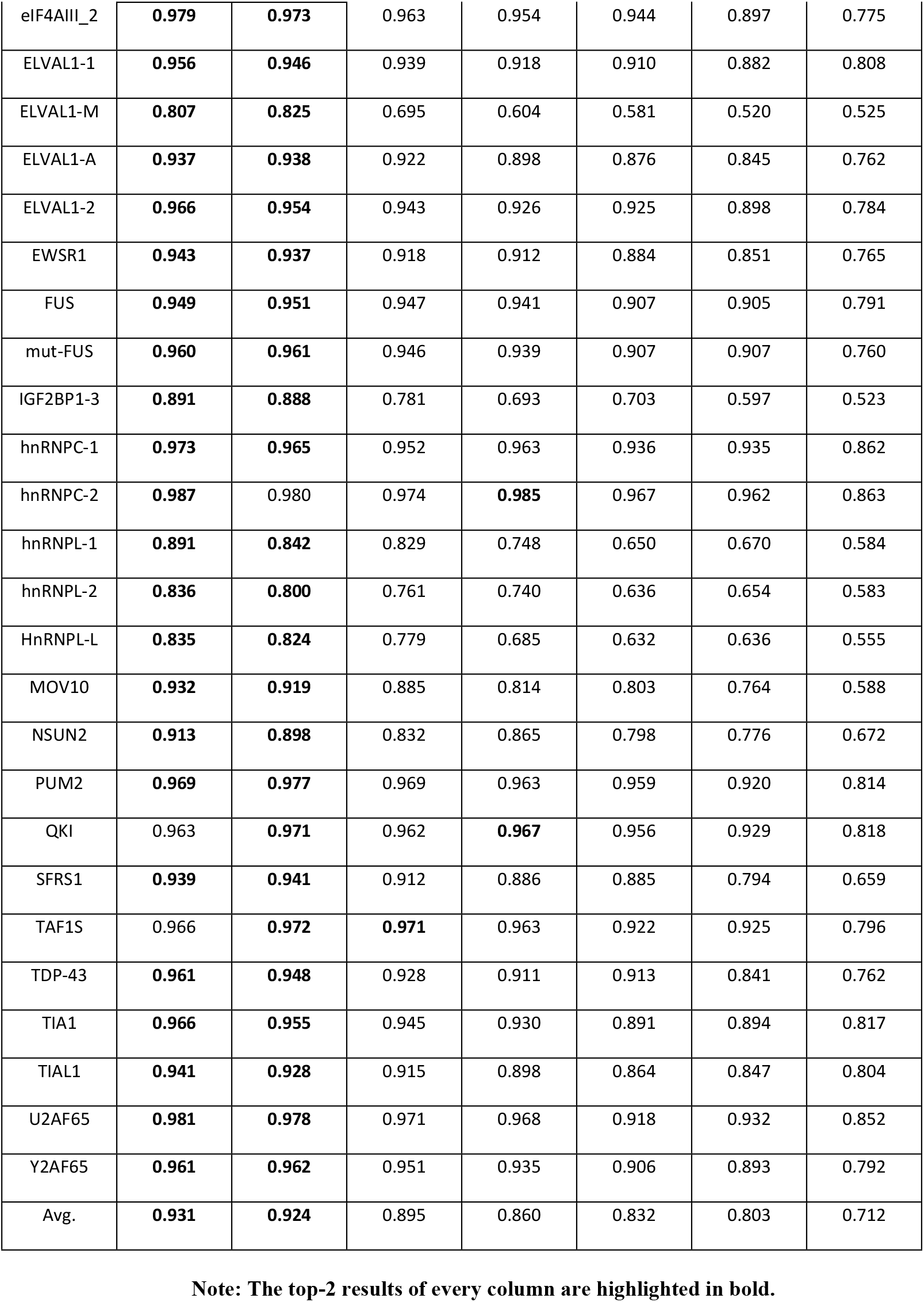
Average value of AUC obtained by different methods on 31 linear RNA datasets.

In addition, to investigate transformability of different methods, we also compared CircSSNN and HCRNet, the newest and most representative algorithm, on linear RNA with their hyper-parameters setting on CircRNA. The experimental results on 31 linear RNA benchmark datasets are shown in Fig 10.

**Fig 10.**
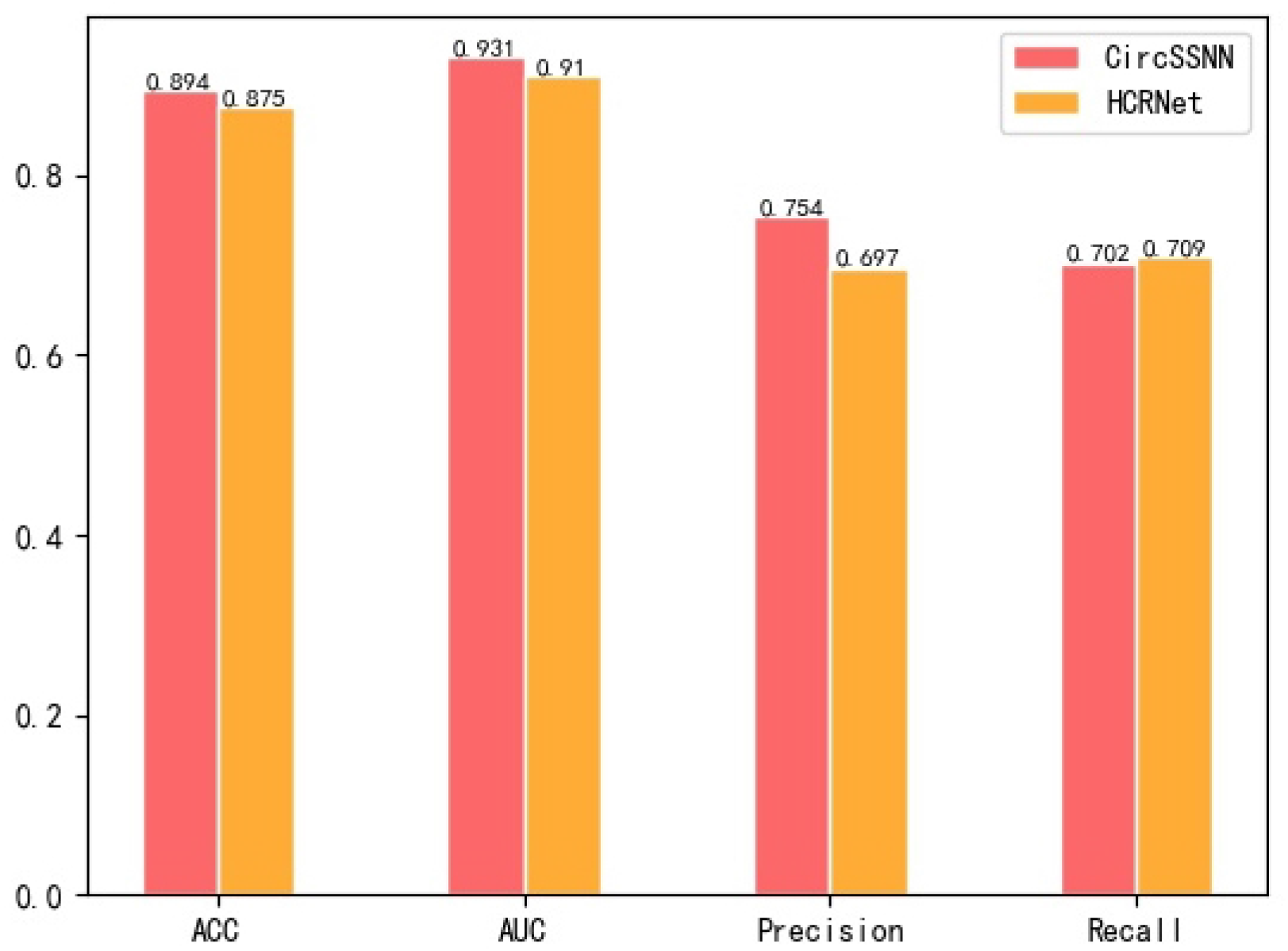
Comparison of CircSSNN and HCRNet on 31 linear RNA datasets.

As shown in Fig 10, when both CircSSNN and HCRNet were tested on linRNA dataset with their hyperparameters settings on the CircRNA, CircSSNN outperformed HCRNet about two, two and six percent regarding ACC, AUC, and Precision, respectively, while just slightly inferior to HCRNet regarding Recall by 0.7 percent. These results verified that CircSSNN was more transformable than HCRNet, and can obtain favorable results even without hyperparameter tuning. The AUC of HCRNet was reported as 0.924 in its original paper, which was the result obtained by fine-tuning the hype-parameters with validate sets, but it dropped to 0.91 when no task-oriented finetuning of hype-parameters is conducted. Therefore, although HCRNet also achieved good performance on the linRNA, the tuning of its hyperparameters requires expertise and a lot of trial and error, which is not conducive to generalization. In contrast, CircSSNN can be simply and efficiently transformed to other RNA-RBP identification tasks and has a wide range of applications.

## Conclusion

At present, most existing models for circRNA-RBP identification adopt CNN, RNN or their variant as feature extractors and have drawbacks such as poor parallelism, insufficient stability, and inability to capture long-term dependence. We propose CircSSNN model based on the sequence self-attention mechanism. CircSSNN extract deep features completely by self-attention mechanism with good parallelism and can capture the long-term dependencies by reducing the distance between any two positions in a sequence to a constant. Multiple experiments on 37 circRNAs and 31 linRNAs datasets using the same hyperparameters show that CircSSNN achieves excellent performance, has good stability and scalability, and eliminates the trouble of hyperparameters tuning compared with existing models. In conclusion, CircSSNN can serve as a appealing option for the task of circRNA-RBP identification.

